# Simulation study of Cell Transmembrane Potential and Electroporation Induced by Time-Varying Magnetic Fields

**DOI:** 10.1101/2022.07.21.501056

**Authors:** Fei Guo, Kun Qian, Xin Li, Hao Deng

**Author notes:** Corresponding author: Fei Guo,.

## Abstract

In this work, an improved simulation model was proposed to assess the transmembrane potential (TMP) evolution on the cellular membrane exposed to time-varying magnetic fields (TMFs). Comparatively, we extended the research on TMP induced by TMF to the electroporation phenomenon by introducing the Smoluchowski function, thereby predicting the occurrence of electroporation. The simulation results based on our numerical model showed that with exposure to the sub-microsecond trapezoidal pulsed magnetic field (PMF), the pore density did not reach the conventional electroporation criterion (10^14^ *m*^−2^) even if the TMP exceeded the electroporation threshold (∼1V); however, with the same energy import, it was easier for the nanosecond pulse to electroporate the membrane evidenced by higher pore density. Further, the capability of predicting the occurrence of electroporation was verified by extending our simulation model to compare experimental results. The comparative analysis showed that our simulation model has predictive and guiding significance for experimental studies and practical applications.

## 1. Introduction

TMFs, including PMFs and oscillating magnetic fields (OMFs), are an emerging non-thermal technique in the food processing industry [1, 2, 3], which can maximize the preservation of food appearance, flavour, texture, and nutritional quality, and improve the freezing process and quality of fresh ground beef [4], chicken breast [5], blueberries [6], cucumber [7], etc. In terms of bactericidal effect, PMF can be used to inhibit and destroy microorganisms such as Escherichia coli [8, 9, 10], yeast [11], and Staphylococcus aureus [12]. Besides, PMF also exhibited a superior anti-bacterial effect on several kinds of bacteria when it is applied in fruit or vegetable juices [13, 14].

Since L. Towhidi *et al*. [15] discovered that the PMF could induce cellular membrane damage and increase its permeability, plenty of empirical studies were commenced to unveil the mechanism of PMF-induced magnetoporation [3, 14, 16, 17, 18, 19, 20], and explore its possibility of applying it to medical practice and food processing. The mechanism of PMF-induced increase in the cellular membrane permeability has been always speculated to be directly related to electroporation (EP) [20, 21, 22]. However, the exact mechanism of PMF-induced EP has not been fully ascertained, which may hinder the application of PMF in the field of food processing. One of the leading hypotheses is that an intense TMF induces a timevarying electric field according to Faraday’s law of electromagnetic induction, which then leads to EP of the cellular membrane [21, 23].

In the meanwhile, simulation studies were conducted and dedicated to revealing the possible micromechanism of the PMF-induced magnetoporation by numerical computation.H. Ye *et al*. [24] calculated the TMP induced by the magnetic field with various frequencies ranging from 2 to 200 Hz in the axon through the uniform cylindrical volume conductor model and analyzed the biophysical properties of the axon as well as the effect of cell polarization. A. Lucinskis *et al*. [22] analyzed the electric field spatial distribution in the cell medium encircled copper wire groups with different structures and inductance. In their simulation work, the TMP of the Jurkat T lymphocyte cells could not reach 0.2 V by analytic computation, inconsistent with reported experimental studies. Q. Hu *et al*. [25] calculated the TMP of spherical cells caused by TMFs and analyzed the influence of various parameter combinations on the time course of the TMP. These simulation studies did not consider the influence of EP on the evolution of TMP, which is of non-negligible importance to the simulation study on the PMF-induced EP. Recently, E. Chiaramello *et al*. [26] focused on the induced TMP by half period sinusoid pulse burst and its dependence on pulse frequency and cell’s position relative to the coil. However, without the verification by comparing to the experimental studies, the theoretical research based on the numerical simulation can barely be instructive to the application of PMF in the practice of experiments and clinics.

Based on the above research, this paper further optimized the simulation settings. The EP effect was introduced to construct the numerical calculation model of cell TMP and pore density under the action of the time-varying pulsed magnetic field. Firstly, the reproduced fundamental model was verified by comparing it to the published numerical studies, and the influence of taking the EP effect into account on the temporal evolution of the TMP was analyzed. Then, based on the improved model, the spatial and temporal evolution of the TMP and pore density was investigated when the cell was exposed to the sub-microsecond and nanosecond pulse. Finally, two sets of experiments were selected for comparative simulation analysis to explore the predictability of experimental results and to verify the instructiveness of understanding the promising application directions of TMFs in the food processing industry.

## 2. Methods

In this study, the fast Fourier transform method (FFT) was applied to transform the time-varying current signal into the Fourier series, thus obtaining harmonic components. Multiplying each term in the Fourier series by the frequency response of TMP to the corresponding frequency, *Ψ*_*tmp*_(*jkω*_0_), the TMP in the time domain in a summation form can be obtained. Introducing the pore density *N*_*p*_(*t*) and the membrane conductivity *σ*_*m*_(*t*), the TMP response taking the EP effect into account 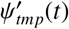 can be finally obtained. The following is the sketchy calculation roadmap, in which the detailed formula derivation and calculation would be expatiated in the following subsections.

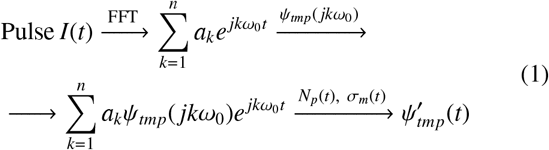

### 2.1. Model details

Fig. 1 shows the three-layer dielectric model of the spherical cell used in this study and its relative spatial position to the coil. The origin of the first Cartesian coordinate system (*X, Y, Z*) is located at the centre of the coil in the *X* − *Y* plane. Viewed from the top or z-axis, a counterclockwise AC current *I*(*t*) was applied to the N-turn loop coil of radius *a*. The centre of the cell, as well as the origin of the second Cartesian coordinate system (*X*′, *Y*′, *Z*′), is located at (0, *c*_*y*_, −*c*_*z*_). The cell was represented as a double-layered sphere with a thin membrane of thickness *d* = *R*^+^ − *R*^−^, and the cellular membrane with inner radius *R*^−^ and outer radius *R*^+^ divided the simulation space into three uniform isotropic regions, the extracellular medium (0#), the cell membrane (1#) and the cytoplasm (2#). It is worth noting that in order to maximize the magnetic vector potential acting on the cellular membrane, thereby prompting the TMP, the cell was placed right under and close to the coil, that is *c*_*y*_ = *a* and *c*_*z*_ = *R*^+^.

**Figure 1:**
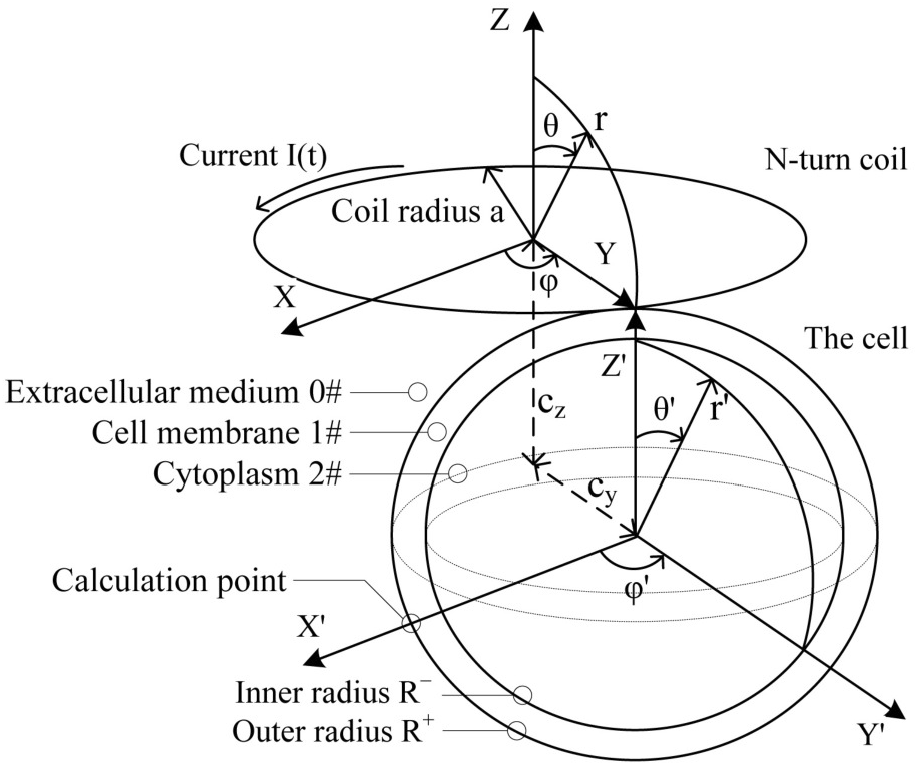
The geometrical model of the cell and its relative position to the coil (Not in scale for details). The centre of the horizontal N-turn coil, whose thickness was ignored, was set to be the origin of the Cartesian coordinate system (*X, Y, Z*), and the centre of the cell located at (0, *c*_*y*_, −*c*_*z*_). The cell was placed right under the coil, which means *c*_*y*_ = *a* and *c*_*z*_ = *R*^+^. The cellular membrane separated the computational domain into three parts, the extracellular medium (0#), the cellular membrane (1#) and the cytoplasm (2#).

### 2.2. Arithmetic of the transmembrane potential

The generated time-varying electromagnetic field changes at the same angular frequency with field sources, the current in this study, vary with time-harmonic quantities (sine or cosine) at a fixed angular frequency [27], and the time-varying field can be expanded as a superposition of time-harmonic fields of different frequencies by Fourier analysis. The induced electric field generated by the time-harmonic electromagnetic field in the biological medium was determined as 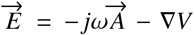 [28], where 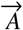 is magnetic vector potential induced by the current in the coil, and *V* is the electric scalar potential induced by the charge accumulation. The boundary condition settings are listed as follows. (i) The normal component of current density 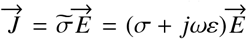 in the boundary (the layers representing the cellular membrane) was continuous across two different media, that is, 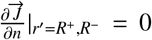. (ii) The scalar electric potential was also continuous across the boundary of two different media.

Transforming the Cartesian coordinate system of the coil into the spherical coordinate system (*r, θ, φ*), the magnetic vector potential components followed 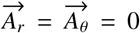. According to the complete elliptic integrals of the first kind (*K*(*m*)) and the second kind (*E*(*m*)), the third component 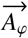 can be expressed as [28]

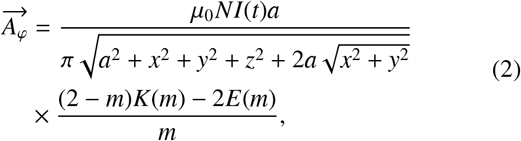

where

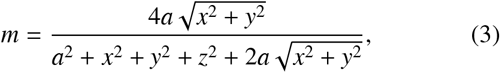

and *µ*_0_ is the vacuum permeability. By space conversion and coordinate transformation, the magnetic vector potential component 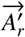 acting on the cell by the coil can be deployed in the spherical coordinates (*r*′, *θ*′, *φ*′)

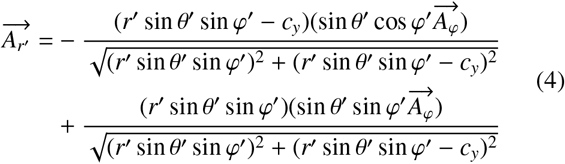

The scalar potential *V* is generated by the charge accumulation of the TMF [29]. The scalar potential of each layer of the medium can be obtained by solving the Laplace equation ∇^2^*V* = 0 in spherical coordinates expression

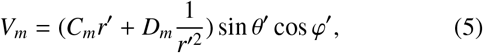

where *C*_*m*_ and *D*_*m*_ are unknown coefficients, and the subscripts *m* = 0, 1, 2 represents the three regions, the extracellular medium (*m* = 0), the cell membrane (*m* = 1) and the cytoplasm (*m* = 2). The potential inside the cell was finite, so it can be obtained by substituting *r*′ = 0 into Eq. 5 that *D*_2_ = 0; the electric field infinitely far away from the cell cannot be interfered with the presence of the cell, so it can be obtained by substitute *r*′ = ∞ into Eq. 5 that *C*_0_ = 0. Across the boundary of two different media, the normal component of the current density and the scalar potential were continuous. Therefore, the parameters *C*_*m*_ and *D*_*m*_ can be solved simultaneously with the above boundary conditions on the outer layer (the boundary between 0# and 1#) and the inner layer (the boundary between 1# and 2#). The response of TMP induced by the *n*-th harmonic of the current signal can be expressed by the scalar potential V on 1# as

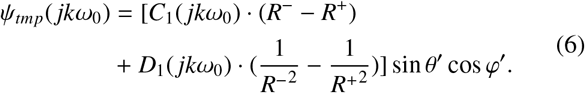

By sampling and FFT analyzing, the current signal on the coil can be expressed in terms of its time-harmonic components 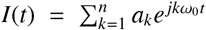. Following the Nyquist Sampling theory and alleviating the calculation load, the number of sampling points was set to be *n* = 2000. By multiplying each time-harmonic component of the current signal by the frequency response of TMP (Eq. 6), the time-domain response of TMP to the applied AC current was obtained by superimposing [30, 31]

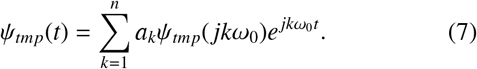

### 2.3. Electroporation effect

EP is a physical phenomenon that occurs on biological membranes. When the potential difference between the two sides of the biofilm reaches a certain threshold, hydrophilic pores are created on the biofilm, resulting in a rapid increase in the conductivity of the biofilm [32]. The occurrence of EP improves membrane permeability instantaneously, thus the mechanism of transmembrane transport of macromolecules can break the normal procedure relying upon the channel proteins. The process of EP includes the formation and development of micropores, which is generally described by Smoluchowski partial differential equation. The main formula used in this paper was the asymptotic model of the EP [33, 34],

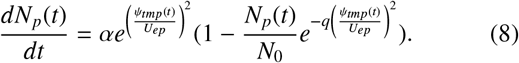

where *U*_*ep*_ is the characteristic voltage of EP, *N*_0_ is the initial pore density of the cell membrane, *α* is the rate of pore formation, and *q* is the EP constant. The above ordinary differential equation was integrated in each time step, from *t*_*n*_ to *t*_*n*+1_, with initial conditions *Ψ*_*tmp*_(*n*) with the stiff solver, ode23t, one of the built-in functions in MATLAB. After EP, highly conductive cytoplasm and extracellular suspension flow into the pores of cellular membrane, resulting in the increase of overall conductivity of cellular membrane

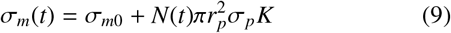

where *σ*_*m*0_ is the static conductivity of the cell membrane, *σ*_*p*_ is the equivalent conductivity of a pore, *r*_*p*_ is the pore radius, and *K* is the partition factor [35].

## 3. Results and discussion

### 3.1. Simulation results in frequency domain and time domain

To test the accuracy of the algorithm in frequency domain (Eq. 6), the corresponding model was built with the dielectric and geometric parameters in Ref. [36]. The frequency spectra of TMP of cellular membrane and organelle membrane were consistent with those in the published study [36]. The comparative results are shown in Fig. 2.

**Figure 2:**
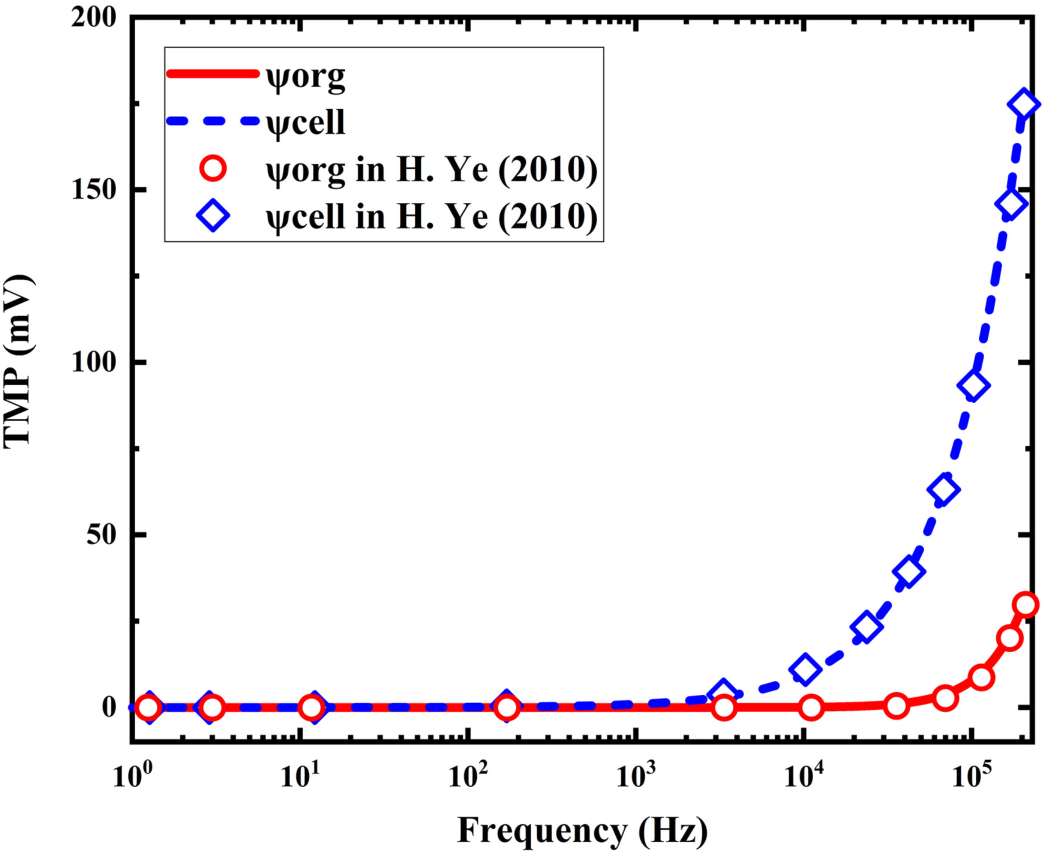
The comparative simulation result of the TMP in the frequency domain at the same probe point as that in [36]. The H. Ye’s TMPs on organelle membrane *Ψ*_*org*_ and cellular membrane *Ψ*_*cell*_ are shown in red circles and blue diamonds, and our simulation results of organelle membrane’s and cellular membrane’s TMPs are shown in red solid line and blue dashed line.

The results showed that the TMP of cellular membrane remained almost constant below 10^3^ Hz, then started to surge rapidly with the increasing frequency, reaching 90 mV at 10^5^ Hz and reaching 170 mV at 2 × 10^5^ Hz. The trend of the TMP of organelle membrane in the frequency spectrum was similar to that of the plasma membrane with a relatively lagging increase with frequency that started at a higher frequency of 10^4^ Hz. For fast input time-varying current, which contains highfrequency components during the rise- and fall-time, it might induce higher TMP of cellular membrane and exceed EP threshold of 1 V [39]. Consistent results were obtained in the simulation verification with Ref. [36]. As the signal frequency increases, TMP shows a tendency to reach higher amplitudes and exceed 1 V at high frequencies.

To verify our algorithm for calculating time-domain TMP induced by time-varying current, the corresponding model was built using the same parameters in Ref. [25]. In the time-domain analysis, the trapezoidal current pulse with 500 A amplitude, 50 ns pulse width and 10 ns rise- and fall-time was applied to the coil, and the non-differentiable waveform was smoothed by the first-order smoothing provided by COMSOL Multiphysics. At the same time, the simulation time was prolonged to reduce the response error at the non-zero signal. The cell was assumed to be directly under and close to the coil, and the TMP calculation point located at *θ*′ = 90°, *φ*′ = 0°, shown in Fig. 1, where the maximum value of TMP on the cell surface can be obtained. The simulation results of TMP and the local pore density in the first 300ns of the simulation time of 1000 ns were shown in Fig. 3.

**Figure 3:**
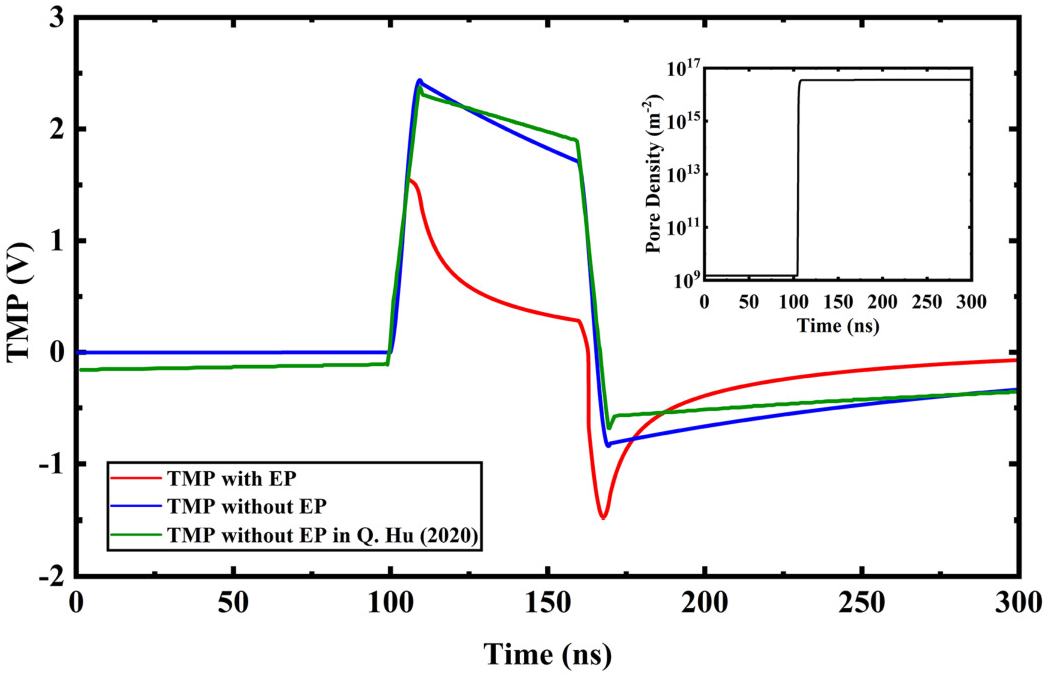
The time-domain TMP simulation results with and without EP effect. The simulation result in [25] was also plotted for comparison. The pore density at the calculation point in the case considering the EP effect was shown in the subgraph.

The results without EP showed that during the pulse rise-time, TMP increased and reached a peak value of 2.44 V, and then gradually decreased to 1.79 V at the end of the pulse flat-top. At the end of the pulse fall-time, TMP decreases to -0.73 V, then after TMP gradually stabilized to nearly 0 V. The overall trend was consistent with the results in Ref. [25], and the possible reasons causing the simulation error are listed as follows: (i) The total simulation time was 1000 ns, longer than that in Ref. [25]. By prolonging the sampling time, which also is the simulation time, the DC component of the signal can be reduced, thereby diminishing its impact on the response of TMP. This procedure can make the TMP before the pulse delivery close to zero. (ii) The pulse applied in our simulation was smoothed by the one-order smoothing, because the idealized trapezoidal pulse is impractical. Meanwhile, the smoothed pulse can reduce the impact of the Gibbs phenomenon on the evolution of TMP [40]. (iii) It is proved in Ref. [25] that in the condition of *c*_*y*_ = *a*, the magnetic potential acting on the cell increases with the lower *c*_*z*_, however, the cell model in the simulation conducted by Hu et al. was not placed close to the coil. Therefore, the subtle difference in cell position may result in the error in Fig. 3.

When the EP effect was introduced, TMP reached its peak of about 1.55 V at 100 ns, then decreased rapidly to 0.34 V at the end of the pulse flattop, further decreased to -1.48 V at the end of the fall-time, finally stabilized to nearly 0 V. The pore density increased rapidly at 100 ns and stabilized to 5 × 10^16^ *m*^−2^. The TMP with EP effect had a relatively obvious downward trend at the flattop time of the pulse in Fig. 3. On the one hand, The change rate of the current amplitude at the flattop of the pulse was zero, then 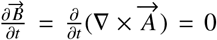, resulting in the absence of the source of the induced electric field. The existing electric field in the computational domain followed the Laplace’s function, therefore, the electric potential difference tended to be zero. On the other hand, with the introduction of the EP effect, the conductivity of cellular membrane increased after EP, which further induced the decrease of TMP. However, the decrease of the TMP was slowed down for the low permittivity of the membrane. After the pulse flattop time, TMP was had been close to zero. During the fall-time of the pulse, the change rate of the magnetic field with respect to time, 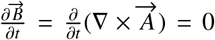, increased negatively, leading to the negative TMP. After the pulse fall-time, the amplitude of TMP with EP effect could recover to close 0 V.

In the comparison simulation with Ref. [25], the TMP evolution without the EP function (Eq. 8 and Eq. 9) obtained by the simulation was basically consistent, but the details were optimized. By extending the simulation time with zero signal, the deviation from zero of the TMP at the non-signal position was reduced. At the same time, by using the first-order smoothed pulse derived from COMSOL Multiphysics, the error caused by the Gibbs phenomenon due to the signal discontinuity was reduced. With the introduction of the EP effect, it can be seen that the peak of the TMP was reduced, the decline of the TMP during the pulse flattop was expedited and the minimum value (negative peak) of the TMP was further reduced.

### 3.2. Sub-microsecond and nanosecond pulse simulation results

The parameters of mouse SP2/0 myeloma cells were used in this subsection, provided in Table. 1 for unified. The sub-microsecond pulse started at 1 *µs* with 0.1 *µs* rise- and fall-time and 1 *µs* half-peak pulse width (plotted in the subgraph in Fig. 4(b)), and during the simulation time of 5 *µs*, the sampling frequency was 0.4 GHz and the time step was 2.5 ns. In the process of gradually increasing the amplitude of pulse, it is observed that the TMP exceeded 1 V and the pore density changed obviously when 1.3 kA pulse current was applied to the 16-turn coil of radius 5 mm. Due to the symmetry of the model, the TMP and pore density on the *X*′+ hemisphere was consistent with the *X*′− hemisphere; because the cell size was much smaller than the coil size, there was little difference between the magnetic potential on the *Y*′+ and *Y*′− hemispheres, also between that on the *Z*′+ and *Z*′− hemispheres, which caused the simulation result between the *Y*′+ and *Y*′− hemispheres and that between the *Z*′+ and *Z*′− hemispheres to be approximately consistent. Therefore, the simulation result of one-eighth of the sphere varying with the azimuth angle *φ*′ and elevation angle *θ*′ was analyzed only, shown in Fig. 4. It can be concluded that on the one-eight of the sphere in the first quadrant of the Cartesian coordinate system (*X*′, *Y*′, *Z*′), the TMP showed a positive relation with the elevation angle *θ*′ when *φ*′ = 0°, and it showed a negative relation with the azimuth angle *φ*′ when *θ*′ = 90°.

**Table 1:** The parameter setting in the model construction.

| Parameter | Value |
| --- | --- |
| Cell inner radius | $5.6 \mu\text{m}$ [37] |
| Membrane thickness | $5\text{nm}$ [37] |
| Membrane permittivity | $4.5 \epsilon_0$ [37] |
| Medium permittivity | $78 \epsilon_0$ [38] |
| Cytoplasm permittivity | $70 \epsilon_0$ [37] |
| Membrane conductivity | $5 \times 10^{-7} \text{ S/m}$ [37] |
| Medium conductivity | $0.17 \text{ S/m}$ [38] |
| Cytoplasm conductivity | $0.25 \text{ S/m}$ [37] |
| Creation rate coefficient ($\alpha$) | $1 \times 10^9 \text{ m}^{-2} \text{ s}^{-1}$ [34] |
| Characteristic voltage of EP ($U_{EP}$) | $258 \text{ mV}$ [34] |
| Equilibrium pore density ($N_0$) | $1.5 \times 10^9 \text{ m}^{-2}$ [34] |
| Pore creation rate ($q$) | $2.46$ [34] |
| Pore radius ($r_p$) | $0.8 \text{ nm}$ [34] |
| Conductivity of a single pore ($\sigma_p$) | $0.22 \text{ S/m}$ [34] |
| Radius of coil ($a$) | $5 \text{ mm}$ [25] |
| Turns of coil ($N$) | $16$ [25] |
| Coil axis-cell distance ($c_y$) | $5 \text{ mm}$ [25] |
| Coil axis-cell distance ($c_z$) | $5.6 \mu\text{m}$ [25] |

**Figure 4:**
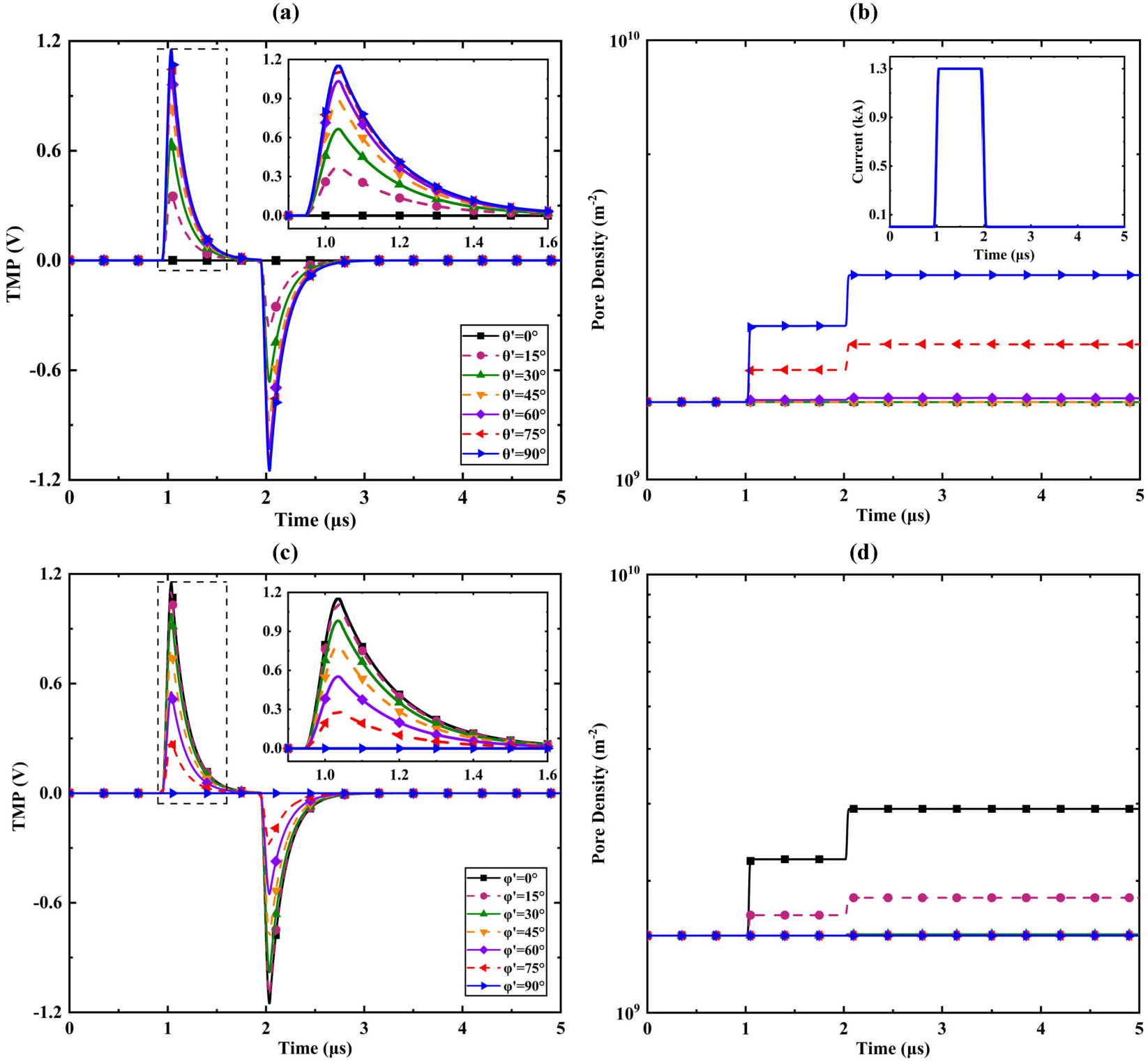
The simulation results of the sub-microsecond pulse case. (a) and (b) show the TMP and pore density varying with elevation angle *θ*′ when the azimuth angle *φ*′ = 0°, that is, the spatio-temporal evolution on the quadrant from the positive *Z*′ axis to the positive *X*′ axis. (c) and (d) show the TMP and pore density varying with azimuth angle *φ*′ when the elevation angle *θ*′ = 90°, namely, those on the quadrant from the positive *X*′ axis to the positive *Y*′ axis. The dashed boxes in (a) and (c) represent the zoom-in area shown in the subgraphs.

Thus, the TMP reached its maximum value at the position of *θ*′ = 90° and *φ*′ = 0°. At this point, TMP increased and reached the peak value of 1.15 V at *t* = 1.04 *µs*, and reached the negative peak value of -1.15 V at *t* = 2.04 *µs*. During the pulse flattop, the TMP could decline to zero due to the long enough pulse flattop. After the cancellation of the pulse, the TMP gradually stabilized to zero, similar to the trend during the pulse flattop. During the rise- and fall-time of the current pulse, the absolute value of TMP exceeded 1 V, and the obvious changes in the pore density were observed. After a two-step increment, the pore density reached 1.5 × 10^10^ *m*^−2^ at this point, though did exceed the the conventional electroporation criterion 10^14^ *m*^−2^ [41, 42].

For further studying the influence of higher frequency pulse on TMP with EP effect, the nanosecond pulse with the same energy was applied to the coil. The nanosecond pulse with 10 ns rise- and fall time and 100 ns half-peak pulse width started at 100 ns. The simulation time in the nanosecond simulation case was 1000 ns, and the sampling frequency was 2 GHz, namely, the time step was 500 ps. The trend of TMP and pore density with azimuth and elevation angle obtained by simulation was shown in Fig. 5. Similar spatial distribution can be observed that the maximum value of TMP occurred at the point of *θ*′ = 90° and *φ*′ = 0°, and TMP decreased with the decreasing *θ*′ or the increasing *φ*′. At this point, the TMP reached the peak of 1.6 V at t=101 ns, and slowly decreased to 0.35 V at t=195 ns. During the fall-time of the pulse, the TMP reached the negative peak of -1.5 V at t=202 ns. The temporal evolution of the pore density was different from that in the sub-microsecond case. Because of the over-intense transient induced electric field, the pore density was saturated and stabilized around 10^16^ *m*^−2^.

**Figure 5:**
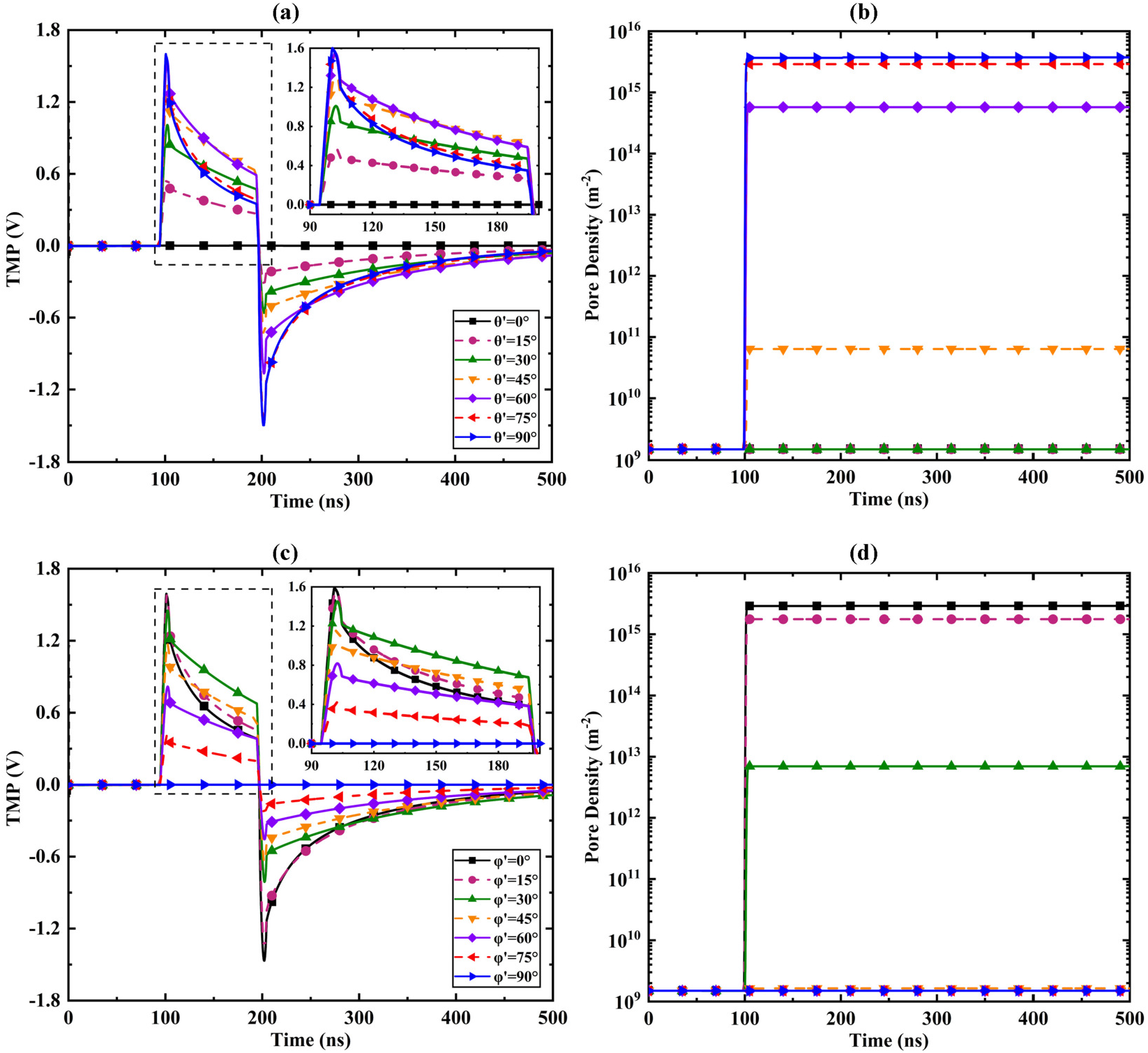
The simulation results of the nanosecond pulse case. (a) and (b) show the TMP and pore density varying with elevation angle *θ*′ when the azimuth angle *φ*′ = 0°. (c) and (d) show the TMP and pore density varying with azimuth angle *φ*′ when the elevation angle *θ*′ = 90°. The dashed boxes in (a) and (c) represent the zoom-in area shown in the subgraphs.

Comparing the above two simulations, it can be inferred that with the same energy import, the higher TMP and pore density can be obtained with a PMF with a higher frequency and a higher amplitude. According to the Faraday law of electromagnetic induction, a TMF with a higher frequency can generate an intenser electric field, leading to the occurrence of EP effect. Similar to the EP studies with pulsed electric fields, in the condition of the equal energy import, the EP can be intensified with a pulse with a higher frequency, supported by the experimental studies [43] (indicated by the shift of fluorescence dyes) and the simulation studies [44] (indicated by the pore density and radius).

### 3.3. Predictability in experimental results

It was reported by the experimental studies that TMFs generated by a multi-turn solenoid can cause reversible EP of cells, increasing membrane permeability, but not affecting cell viability [45, 46]. Two pieces of experimental research were chosen to verify the predictability of our simulation model.

The first experimental research conducted by V. Novickij et al. [47] used a solenoid type inductor with a 16-winding coil, stacked with two layers of the 8-winding coil, and placed Jurkat T-lymphocytes in the inner container of the coil. Comparing the number of fluorescent cells in the experimental group and the control group, it showed that the EP could not be induced with the pulse parameter setting in the study. The parameter setting to fit the simulation research was detailed in Table. 2. The damped sinusoidal signal used in Ref. [47] and the fitting signal used in our simulation were plotted in Fig. 6(a). The error between the experimental signal and the idealized simulation signal was ignored because it is hard to perfectly fit the practical signal with the fundamental mathematical functions and the error during the low amplitude period can hardly affect the EP degree. Fig. 6(b) shows the maximum value on the cellular membrane and the corresponding pore density calculated by our model. TMP oscillated damply following the fluctuating frequency of current signal, and during the simulation time period, the peak of TMP reached only 0.7 V, not exceeding the electroporation threshold of 1 V. Accordingly, the pore density remained at its initial value, indicating that the EP effect did not occur. Therefore, the simulation result proved that this parameter setting cannot meet the conditions for EP occurring, consisting of the experimental result.

**Table 2:** The parameter setting in predicting the experimental results.

| Parameter Type | Parameter | Value for [47] | Value for [48] |
| --- | --- | --- | --- |
| Cell parameters | Cell radius | $7 \mu\text{m}$ [49] | $5.6 \mu\text{m}$ [37] |
| | Membrane thickness | $5.9 \text{ nm}$ [49] | $5 \text{ nm}$ [37] |
| | Membrane permittivity | $5.8 \epsilon_0$ [49] | $4.5 \epsilon_0$ [37] |
| | Medium permittivity | $78 \epsilon_0$ [38] | $78 \epsilon_0$ [38] |
| | Cytoplasm permittivity | $60 \epsilon_0$ [49] | $70 \epsilon_0$ [37] |
| | Membrane conductivity | $8.7 \times 10^{-6} \text{ S/m}$ [49] | $5 \times 10^{-7} \text{ S/m}$ [37] |
| | Medium conductivity | $0.17 \text{ S/m}$ [38] | $0.17 \text{ S/m}$ [38] |
| | Cytoplasm conductivity | $0.48 \text{ S/m}$ [49] | $0.25 \text{ S/m}$ [37] |
| Coil parameters | Radius of coil($a$) | $0.75 \text{ mm}$ [47] | $5.2 \text{ mm}$ [48] |
| | Turns of coil($N$) | 16 [47] | 6 [48] |
| | Coil axis-cell distance ($c_y$) | $7 \mu\text{m}$ [47] | $5.6 \mu\text{m}$ [48] |
| | Coil axis-cell distance ($c_z$) | $5.9 \text{ nm}$ [47] | $5 \text{ nm}$ [48] |
|  | Current peak value | 1200 A [47] | 5000 A [48] |
| Damped sinusoidal signal parameters<br>$Ae^{-\xi t} \sin(\omega_n t + \varphi_n)$ | Amplitude($A$) | -1.079 | -1.069 |
| | Damping ratio($\xi$) | $0.208 \times 10^6$ | $0.179 \times 10^6$ |
| | Frequency($\omega_n$) | $1.352\pi \times 10^6$ | $1.428\pi \times 10^6$ |
| | Phase angle($\varphi_n$) | $-\pi$ | $-\pi$ |
| Simulation settings | Time step | 10 ns | 14 ns |
| | Simulation time | 6 $\mu\text{s}$ | 14 $\mu\text{s}$ |
|  | Sampling frequency | 100 MHz | 70 MHz |

**Figure 6:**
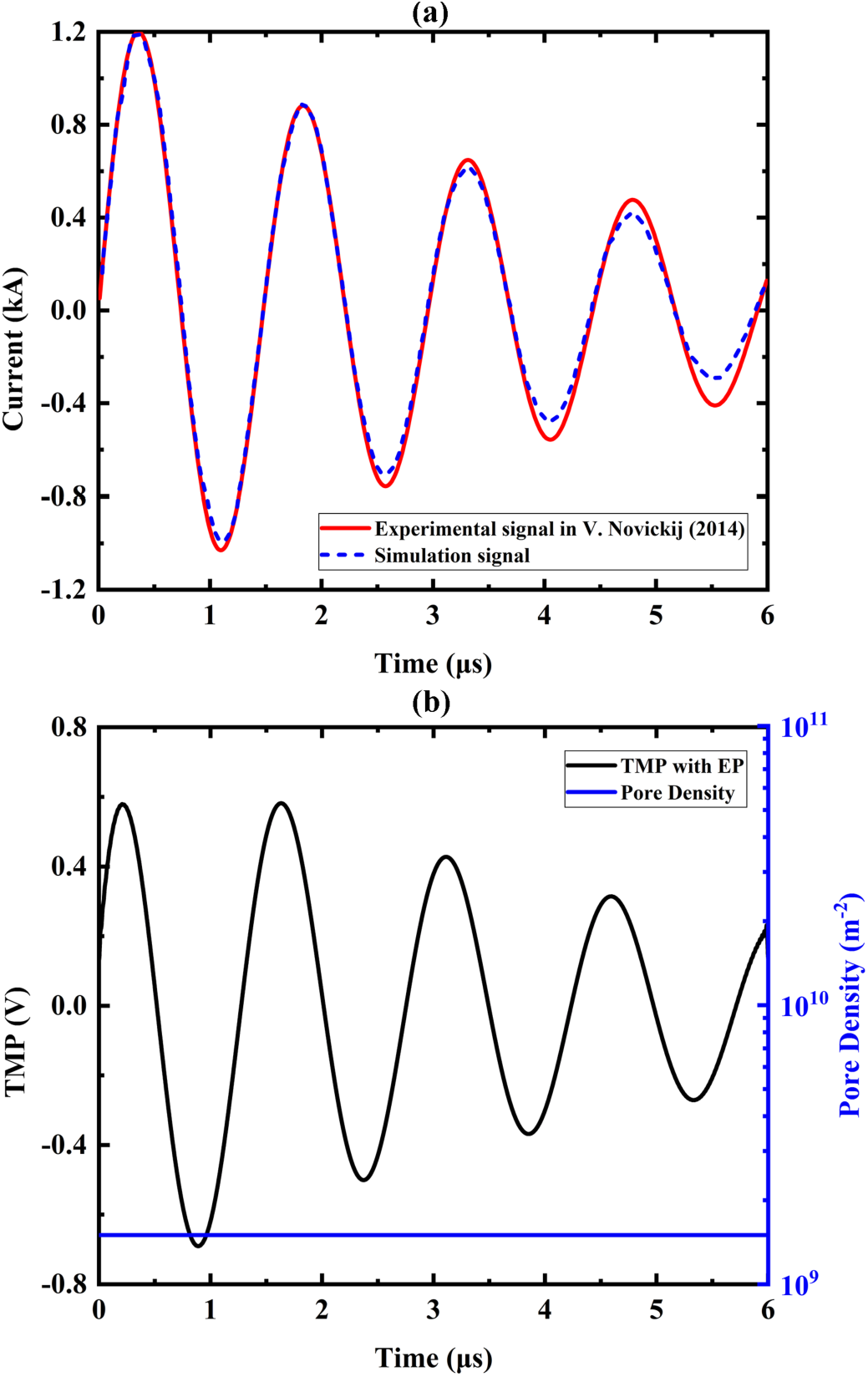
Comparison between the experimental research conducted by V. Novickij et al. [47] and the simulation result from our model. (a) The damped sinusoidal current signal applied to the coil. (b) the maximum value of TMP on the cell and the corresponding pore density.

The other experimental research referred to verify the predictability of our model was also conducted by V. Novickij et al. [48]. A solenoid-type inductor with a 6-winding coil stacked with two layers of the 3-winding coil was applied to import the current signal, and the SP2/0 myeloma cell was placed in the inner container of the coil. Accordingly, the simulation model was built with the corresponding parameter setting, also detailed in Table. 2. The damped sinusoidal current signal recorded in the experiment was shown in Fig. 7(a). By analyzing the shift of YO-PRO-1 fluorescent, the EP degree indicated by the membrane permeability can be estimated. With the fitting current signal applied in our model, the maximum value of TMP calculated on the membrane was shown in Fig. 7(b). When TMP reached the first positive peak of 0.98 V at *t* = 1.6 *µs*, the pore density made the first small step. When the TMP dropped over -1 V during the decline in the first period of the sinusoidal signal, the pore density surged to 10^13^ *m*^−2^, indicating the occurrence of the EP effect.

**Figure 7:**
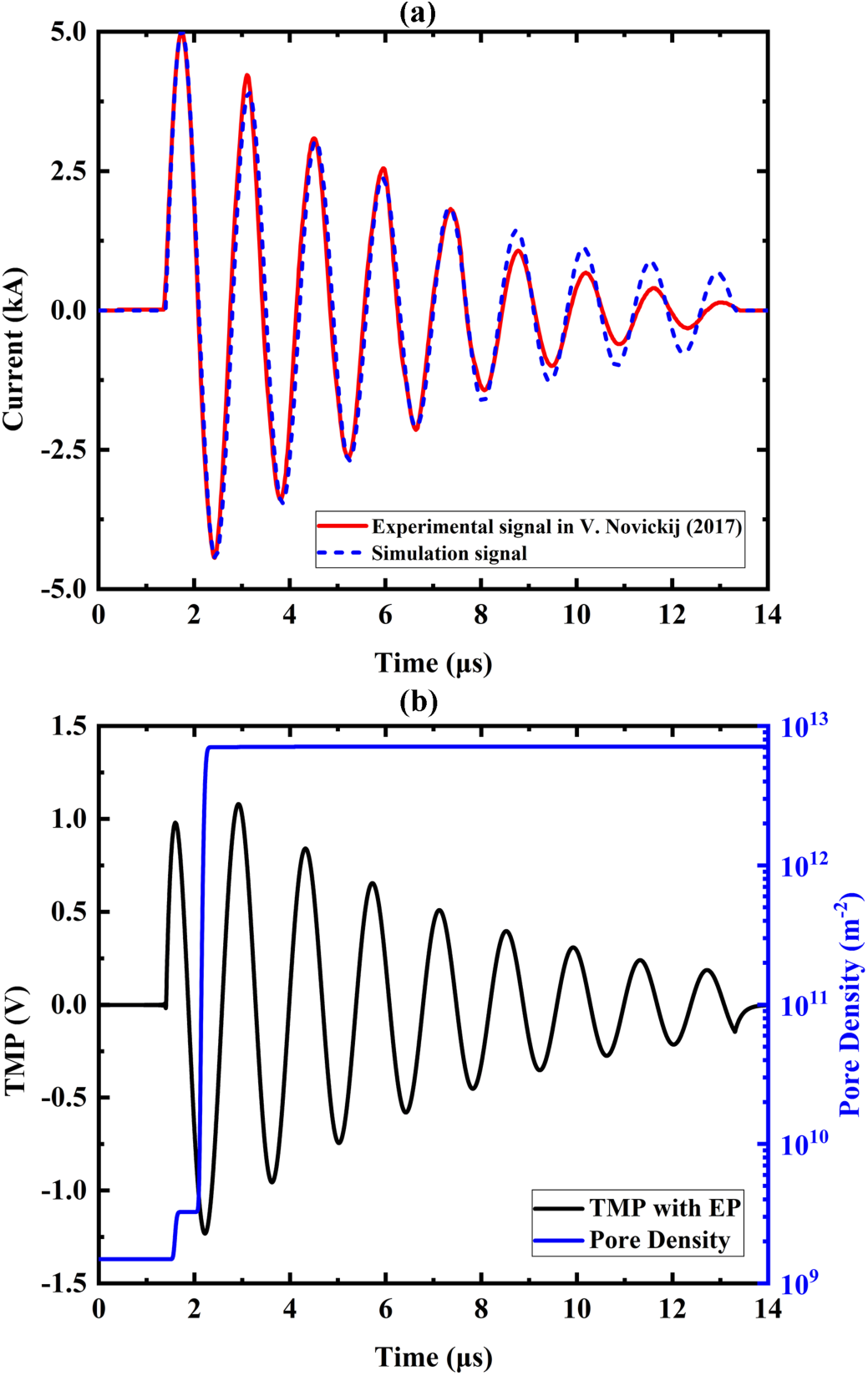
Comparison between the second experimental research also conducted by V. Novickij et al. [48] and the simulation result from our model. (a) The damped sinusoidal current signal applied to the coil. (b) the maximum value of TMP on the cell and the corresponding pore density.

In comparison with the experimental results of Ref. [47], the TMP did not reach the commonly considered electroporation threshold, and the pore density remained at its initial value, so it can be considered that no electroporation phenomenon occurred, consistent with the experimental result. In comparison with the experimental results of Ref. [48], the TMP exceeded the threshold, and the pore density rose to around 10^13^ *m*^−2^, lower than the generally considered pore density threshold of 10^14^ *m*^−2^. It ascribes the deviation to the difference in parameter settings because it is hard to obtain the accurate parameters in the compared experiment studies, such as the conductivity and permittivity of the medium even if the parameters of the same cell line were selected and applied in this study. In addition, as mentioned before, the mainstream hypothesis for magnetoporation is that the PMF induces the time-varying electric field, and as the conventional electric field induced electroporation, the electric field acts on the cellular membrane inducing TMP, thereby breaking the isolation effect of the cellular membrane. However, experimental research indicated that estimated PMF-induced electric field strength is much lower from those commonly used for conventional electroporation, but still increases the membrane permeability *in vivo* and *in vitro* [20, 26, 46], implying that other aspects introduced by PMF influence the magnetoporation. Hydrostatic pressure induced by PMF could be one of the influencing factors. Hydrostatic pressure could not only affect the pores opening energetic balance by activating or deactivating the ion channels, but also contributes additional redial pressure on cellular membrane, enhancing the probability of opening pores [26, 50, 51]. Another influencing factor could be the inevitable Joule heating of the inductors due to the high current in the coils [46, 52]. The temperature rise when the high currents up to several kA in the coil would significantly affects the pore forming process and the vitality of cells/microorganisms [53, 54]. In general, it is observed that considerably lower PMF-induced electric field strength demanded for membrane permeabilization compared to the conventional electroporation process. The underlying mechanism is still opaque, requiring further investigation in future research.

## 4. Conclusion

TMF technology offers an efficient physical sterilization protocol for food processing industry and a contactless and noninvasive method for cancer treatment. A multiphysics model, based on the hypothesis that PMF-induced electric field results in membrane electroporation, was proposed to preliminarily assess the TMP during the PMF delivery and estimate the pore forming process on the cellular membrane. This model has guiding significance for the coil design and the selection of current signal parameters in subsequent experiments. To extent the simulation model to future research, the mentioned biomechanics on a micro scale and temperature rise could be included to contribute to a profound understanding of the underlying mechanism of PMF-induced cellular membrane permeabilization.

## Declaration of Competing Interest

The authors declare that they have no known competing financial interests or personal relationships that could have appeared to influence the work reported in this paper.

## Acknowledgement

This work was supported in part by the Science and Technology Research Program of Chongqing Municipal Education Commission (Grant No.KJQN202100607), in part by the Natural Science Foundation of Chongqing, China (cstc2020jcyj-msxmX0393), and in part by the National Natural Science Foundation of China (No. 51507024).

## Notes

### Competing Interest Statement

The authors have declared no competing interest.

## References

[1] A. M. El-Khatib, A. M. Khalil, M. I. El-Kaliuoby, M. Elkhatib, The combined effects of multisized silver nanoparticles and pulsed magnetic field on k. pneumoniae, Bioinspired, Biomimetic and Nanobiomaterials 8 (2) (2019) 154–160. doi:10.1680/jbibn.18.00042.

[2] C. Piyadasa, T. R. Yeager, S. R. Gray, M. B. Stewart, H. F. Ridgway, C. Pelekani, J. D. Orbell, Antimicrobial effects of pulsed electromagnetic fields from commercially available water treatment devices – controlled studies under static and flow conditions, Journal of Chemical Technology & Biotechnology 93 (3) (2018) 871–877. doi:https://doi.org/10.1002/jctb.5442.

[3] P. Wu, W. Qu, M. A. Y. Abdualrahman, Y. Guo, K. Xu, H. Ma, Study on inactivation mechanisms of listeria grayi affected by pulse magnetic field via morphological structure, ca2+ transmembrane transport and proteomic analysis, International Journal of Food Science & Technology 52 (9) (2017) 2049–2057. doi:https://doi.org/10.1111/ijfs.13483.

[4] P. Goldschmidt Lins, A. Aparecida Silva, S. Marina Piccoli Pugine, A. Ivan Cespedes Arce, E. José Xavier Costa, M. Pires De Melo, Effect of exposure to pulsed magnetic field on microbiological quality, color and oxidative stability of fresh ground beef, Journal of Food Process Engineering 40 (2) (2017) e12405. doi:https://doi.org/10.1111/jfpe.12405.

[5] J. H. Mok, J.-Y. Her, T. Kang, R. Hoptowit, S. Jun, Effects of pulsed electric field (pef) and oscillating magnetic field (omf) combination technology on the extension of supercooling for chicken breasts, Journal of Food Engineering 196 (2017) 27–35. doi:https://doi.org/10.1016/j.jfoodeng.2016.10.002.

[6] J. Tang, S. Shao, C. Tian, Effects of the magnetic field on the freezing process of blueberry, International Journal of Refrigeration 113 (2020) 288–295. doi:https://doi.org/10.1016/j.ijrefrig.2019.12.022.

[7] L. Zhang, Z. Yang, S. Zhao, N. Luo, Q. Deng, Effect of combined pulsed magnetic field and cold water shock treatment on the preservation of cucumbers during postharvest storage, Food and Bioprocess Technology 13 (4) (2020) 732–738. doi:https://doi.org/10.1007/s11947-020-02425-w.

[8] R. He, H. Ma, H. Wang, Inactivation of e. coli by high-intensity pulsed electromagnetic field with a change in the intracellular ca2+ concentration, Journal of Electromagnetic Waves and Applications 28 (4) (2014) 459–469. doi:10.1080/09205071.2013.874539.

[9] L. Lin, X. Wang, H. Cui, Synergistic efficacy of pulsed magnetic fields and litseacubeba essential oil treatment against escherichia coli o157:h7 in vegetable juices, Food Control 106 (2019) 106686. doi:10.1016/j.foodcont.2019.06.012.

[10] V. Novickij, R. Stanevičienė, R. Gruškienė, K. Badokas, J. Lukša, J. Sereikaitė, K. Mažeika, N. Višniakov, J. Novickij, E. Servienė, Inactivation of bacteria using bioactive nanoparticles and alternating magnetic fields, Nanomaterials 11 (2) (2021). doi:10.3390/nano11020342.

[11] S. Mercado-Sáenz, B. López-Díaz, A. M. Burgos-Molina, F. Sendra-Portero, A. González-Vidal, M. J. Ruiz-Gómez, Exposure of s. cerevisiae to pulsed magnetic field during chronological aging could induce genomic dna damage, International Journal of Environmental Health Research 0 (0) (2021) 1–12, pMID: 33797308. doi:10.1080/09603123.2021.1910212.

[12] S. K. Boda, K. Ravikumar, D. K. Saini, B. Basu, Differential viability response of prokaryotes and eukaryotes to high strength pulsed magnetic stimuli, Bioelectrochemistry 106 (2015) 276–289. doi:10.1016/j.bioelechem.2015.07.009.

[13] J. Qian, G. Yan, S. Huo, C. Dai, H. Ma, J. Kan, Effects of pulsed magnetic field on microbial and enzymic inactivation and quality attributes of orange juice, Journal of Food Processing and Preservation 45 (6) (2021) e15533. doi:10.1111/jfpp.15533.

[14] L. Lin, X. Wang, R. He, H. Cui, Action mechanism of pulsed magnetic field against e. coli o157:h7 and its application in vegetable juice, Food Control 95 (2019) 150–156. doi:10.1016/j.foodcont.2018.08.011.

[15] L. Towhidi, S. Firoozabadi, H. Mozdarani, D. Miklavcic, Lucifer yellow uptake by cho cells exposed to magnetic and electric pulses, Radiology and Oncology 46 (2) (2012) 119–125. doi:10.2478/v10019-012-0014-2.

[16] Z. Shankayi, S. M. P. Firoozabadi, M. G. Mansurian, The effect of pulsed magnetic field on the molecular uptake and medium conductivity of leukemia cell, Cell Biochemistry and Biophysics 65 (2013) 211–216. doi:10.1007/s12013-012-9422-6.

[17] S. Lakshmanan, G. K. Gupta, P. Avci, R. Chandran, M. Sadasivam, A. E. S. Jorge, M. R. Hamblin, Physical energy for drug delivery; poration, concentration and activation, Advanced Drug Delivery Reviews 71 (2014) 98–114. doi:10.1016/j.addr.2013.05.010.

[18] V. Novickij, A. Grainys, I. Kučinskaitė-Kodzė, A. Žvirblienė, J. Novickij, Magneto-permeabilization of viable cell membrane using high pulsed magnetic field, IEEE Transactions on Magnetics 51 (9) (2015) 1–5. doi:10.1109/TMAG.2015.2439638.

[19] J. Qian, C. Zhou, H. Ma, S. Li, A. E. A. Yagoub, M. A. Y. Abdualrahman, Biological effect and inactivation mechanism of bacillus subtilis exposed to pulsed magnetic field: Morphology, membrane permeability and intracellular contents, Food Biophysics 11 (4) (2016) 429–435. doi:10.1007/s11483-016-9442-7.

[20] D. Miklavcic, V. Novickij, M. Kranjc, T. Polajzer, S. Haberl Meglic, T. Batista Napotnik, R. Romih, D. Lisjak, Contactless electroporation induced by high intensity pulsed electromagnetic fields via distributed nanoelectrodes, Bioelectrochemistry 132 (2020) 107440. doi:10.1016/j.bioelechem.2019.107440.

[21] T. J. Kardos, D. P. Rabussay, Contactless magneto-permeabilization for intracellular plasmid dna delivery in-vivo, Human Vaccines & Immunotherapeutics 8 (11) (2012) 1707–1713. doi:10.4161/hv.21576.

[22] A. Lucinskis, V. Novickij, A. Grainys, J. Novickij, S. Tolvaisiene, Modelling the cell transmembrane potential dependence on the structure of the pulsed magnetic field coils, Elektronika ir Elektrotechnika 20 (8) (2014) 9–12. doi:10.5755/j01.eee.20.8.8432.

[23] V. Novickij, J. Dermol, A. Grainys, M. Kranjc, D. Miklavčič, Membrane permeabilization of mammalian cells using bursts of high magnetic field pulses, PeerJ 5 (2017) e3267. doi:10.7717/peerj.3267.

[24] H. Ye, M. Cotic, M. G. Fehlings, P. L. Carlen, Transmembrane potential generated by a magnetically induced transverse electric field in a cylindrical axonal model, Medical & Biological Engineering & Computing 49 (1) (2011) 107–119. doi:10.1007/s11517-010-0704-0.

[25] Q. Hu, R. P. Joshi, D. Miklavčič, Calculations of cell transmembrane voltage induced by time-varying magnetic fields, IEEE Transactions on Plasma Science 48 (4) (2020) 1088–1095. doi:10.1109/TPS.2020.2975421.

[26] E. Chiaramello, S. Fiocchi, M. Bonato, S. Gallucci, M. Benini, M. Parazzini, Cell transmembrane potential in contactless permeabilization by timevarying magnetic fields, Computers in Biology and Medicine 135 (2021) 104587. doi:10.1016/j.compbiomed.2021.104587.

[27] D. K. Cheng, et al., Field and wave electromagnetics, Pearson Education India, 1989.

[28] J. D. Jackson, Classical electrodynamics, John Wiley & Sons, 1998.

[29] J. A. Stratton, Electromagnetic theory, Vol. 33, John Wiley & Sons, 2007.

[30] V. K. Ingle, J. G. Proakis, Digital signal processing using matlab: a problem solving companion, Cengage Learning, 2016.

[31] A. V. Oppenheim, A. S. Willsky, S. H. Nawab, G. M. Hernández, et al., Signals & systems, Pearson Educación, 1997.

[32] L. Mittal, I. G. Camarillo, U. K. Aryal, R. Sundararajan, Global proteomic analysis of breast cancer cell plasma membrane electroporation, in: 2019 IEEE Conference on Electrical Insulation and Dielectric Phenomena (CEIDP), 2019, pp. 725–728. doi:10.1109/CEIDP47102.2019.9009762.

[33] J. C. Neu, W. Krassowska, Asymptotic model of electroporation, Phys. Rev. E 59 (1999) 3471–3482. doi:10.1103/PhysRevE.59.3471.

[34] C. Yao, H. Liu, Y. Zhao, Y. Mi, S. Dong, Y. Lv, Analysis of dynamic processes in single-cell electroporation and their effects on parameter selection based on the finite-element model, IEEE Transactions on Plasma Science 45 (5) (2017) 889–900. doi:10.1109/TPS.2017.2681433.

[35] L. Chernomordik, S. Sukharev, S. Popov, V. Pastushenko, A. Sokirko, I. Abidor, Y. Chizmadzhev, The electrical breakdown of cell and lipid membranes: the similarity of phenomenologies, Biochimica et Biophysica Acta (BBA) - Biomembranes 902 (3) (1987) 360–373. doi:10.1016/0005-2736(87)90204-5.

[36] H. Ye, M. Cotic, E. E. Kang, M. G. Fehlings, P. L. Carlen, Transmembrane potential induced on the internal organelle by a time-varying magnetic field: a model study, Journal of NeuroEngineering and Rehabilitation 7 (1) (2010) 12. doi:10.1186/1743-0003-7-12.

[37] C. Li, Q. Ke, C. Yao, C. Yao, Y. Mi, M. Wu, L. Ge, Comparison of bipolar and unipolar pulses in cell electrofusion: Simulation and experimental research, IEEE Transactions on Biomedical Engineering 66 (5) (2019) 1353–1360. doi:10.1109/TBME.2018.2872909.

[38] E. Salimi, K. Braasch, M. Butler, D. J. Thomson, G. E. Bridges, Dielectrophoresis study of temporal change in internal conductivity of single cho cells after electroporation by pulsed electric fields, Biomicrofluidics 11 (1) (2017) 014111. doi:10.1063/1.4975978.

[39] W. K. Neu, J. C. Neu, Theory of Electroporation, Springer US, Boston, MA, 2009, pp. 133–161. doi:10.1007/978-0-387-79403-7_7.

[40] D. Gottlieb, C.-W. Shu, On the gibbs phenomenon and its resolution, SIAM Review 39 (4) (1997) 644–668. doi:10.1137/S0036144596301390.

[41] L. Retelj, G. Pucihar, D. Miklavčič, Electroporation of intracellular liposomes using nanosecond electric pulses—a theoretical study, IEEE Transactions on Biomedical Engineering 60 (9) (2013) 2624–2635. doi: 10.1109/TBME.2013.2262177.

[42] F. Guo, K. Qian, L. Zhang, H. Deng, X. Li, J. Zhou, J. Wang, Anisotropic conductivity for single-cell electroporation simulation with tangentially dispersive membrane, Electrochimica Acta 385 (2021) 138426. doi: 10.1016/j.electacta.2021.138426.

[43] D. Navickaite, P. Ruzgys, V. Novickij, M. Jakutaviciute, M. Maciulevicius, R. Sinceviciute, S. Satkauskas, Extracellular-ca2+-induced decrease in small molecule electrotransfer efficiency: Comparison between microsecond and nanosecond electric pulses, Pharmaceutics 12 (5) (2020). doi:10.3390/pharmaceutics12050422.

[44] F. Guo, K. Qian, L. Zhang, X. Liu, H. Peng, Multiphysics modelling of electroporation under uni- or bipolar nanosecond pulse sequences, Bioelectrochemistry 141 (2021) 107878. doi:10.1016/j.bioelechem.2021.107878.

[45] V. Novickij, J. Dermol, A. Grainys, M. Kranjc, D. Miklavčič, Membrane permeabilization of mammalian cells using bursts of high magnetic field pulses, PeerJ 5 (2017) e3267. doi:10.7717/peerj.3267.

[46] V. Novickij, A. Grainys, E. Lastauskienė, R. Kananavičiūtė, D. Pamedytytė, L. Kalėdienė, J. Novickij, D. Miklavčič, Pulsed electromagnetic field assisted in vitro electroporation: A pilot study, Scientific Reports 6 (1) (2016) 33537. doi:10.1038/srep33537.

[47] V. Novickij, A. Grainys, J. Novickij, Contactless dielectrophoretic manipulation of biological cells using pulsed magnetic fields, IET Nanobiotechnology 8 (2) (2014) 118–122. doi:https://doi.org/10.1049/iet-nbt.2012.0039.

[48] V. Novickij, I. Girkontaitė, A. Zinkevičienė, J. Švedienė, E. Lastauskienė, A. Paškevičius, S. Markovskaja, J. Novickij, Reversible permeabilization of cancer cells by high sub-microsecond magnetic field, IEEE Transactions on Magnetics 53 (11) (2017) 1–4. doi:10.1109/TMAG.2017.2719699.

[49] I. Ermolina, Y. Polevaya, Y. Feldman, B.-Z. Ginzburg, M. Schlesinger, Study of normal and malignant white blood cells by time domain dielectric spectroscopy, IEEE Transactions on Dielectrics and Electrical Insulation 8 (2) (2001) 253–261. doi:10.1109/94.919948.

[50] T. Polyakova, V. Zablotskii, A. Dejneka, Cell membrane pore formation and change in ion channel activity in high-gradient magnetic fields, IEEE Magnetics Letters 8 (2017) 1–5. doi:10.1109/LMAG.2017.2732361.

[51] H. Ye, A. Curcuru, Biomechanics of cell membrane under low-frequency time-varying magnetic field: a shell model, Medical & Biological Engineering & Computing 54 (12) (2016) 1871–1881. doi:10.1007/s11517-016-1478-9.

[52] S. Bartkevicius, J. Novickij, The influence of pulsed magnet heating on maximal value of generated magnetic field, Measurement Science Review 8 (4) (2008) 94. doi:10.2478/v10048-008-0022-y.

[53] V. Novickij, A. Grainys, J. Švedienė, S. Markovskaja, J. Novickij, Joule heating influence on the vitality of fungi in pulsed magnetic fields during magnetic permeabilization, Journal of Thermal Analysis and Calorimetry 118 (2) (2014) 681–686. doi:10.1007/s10973-014-3735-1.

[54] Z. Yan, C. Hao, L. Yin, K. Liu, J. Qiu, Simulation of the influence of temperature on the dynamic process of electroporation based on finite element analysis, IEEE Transactions on Plasma Science 49 (9) (2021) 2839–2850. doi:10.1109/TPS.2021.3100878.

